# Interpretable Multimodality Embedding Of Cerebral Cortex Using Attention Graph Network For Identifying Bipolar Disorder

**DOI:** 10.1101/671339

**Authors:** Huzheng Yang, Xiaoxiao Li, Yifan Wu, Siyi Li, Su Lu, James S. Duncan, James C. Gee, Shi Gu

## Abstract

Bipolar Disorder (BP) is a mental disorder that affects 1 ∼ 2% of the population. Early diagnosis and targeted treatment can benefit from associated biological markers. The existing methods typically utilize biomarkers from anatomical MRI or functional BOLD imaging, but lack the ability of revealing the relationship between integrated modalities and disease. In this paper, we developed an Edge-weighted Graph Attention Network (EGAT) with Dense Hierarchical Pooling (DHP), to better understand the underlying roots of the disorder from the view of structure-function integration. For the input, the underlying graphs are constructed from functional connectivity matrices and the nodal features consist of both the anatomical features and the statistics of the connectivity. We investigated the potential benefits of using EGAT to classify BP vs. Healthy Control (HC). Compared with traditional machine learning classifiers, our proposed EGAT embedding increased improved 10 ∼ 20% in the accuracy and F1-score, compared with alternative classifiers. More specifically, by examining the attention map and gradient sensitivity of nodal features, we indicated that associated with the abnormality of anatomical geometric properties, multiple interactive patterns among Default Mode, Fronto-parietal and Cingulo-opercular networks contribute to identifying BP.

## 1 Introduction

Bipolar disorder (BP), formerly called manic depression, is a mental health condition that causes extreme mood swings that include emotional highs (mania or hypomania) and lows (depression) [1]. Despite decades of research, the pathophysiology of BP is still not well understood. Some of the most commonly prescribed presentation for patients with BP have also been associated with structural or functional brain differences. For example, [8] found adults with BP had widespread bilateral patterns of reduced cortical thickness in the frontal, temporal and parietal regions. Some studies have also shown evidence of reductions in functional connectivity within the cortical control networks [2,22].

Many brain imaging techniques including functional MRI (fMRI), structural MRI (sMRI), EEG/MEG and diffusion tensor imaging (DTI) provide information on different aspects of the brain. Although functional and structural brain studies have identified quantitative differences between BP and Health Control (HC) groups, most models favor only one data type or do not combine data from different imaging modalities effectively, thus missing potentially important differences which are only partially detected by single modality [4,3]. Combining modalities may thus uncover previously hidden relationships that can unify disparate findings in neuroimaging. To the best of our knowledge, no previous work has been done to combine structural and functional connectivity data to analyze BP. We hold the hypothesis that with joint information, the better representation can be learned to describe BPs’ characteristics, and validate this hypothesis in our experiment. A main challenge in multimodal data fusion comes from the dissimilarity of the data types being fused and result interpretation. Traditional multi-modality studies on neuroimaging mainly use principal component analysis (PCA), independent component analysis (ICA), canonical correlation analysis (CCA), and partial least squares (PLS) [19]. However, the model’s intrinsic dependence on the shape and scale of the data distribution causes ambiguity in components discovery and harms the easiness of interpretation.

Graph-based approach for multi-modality is a powerful technique to characterize the architecture of human brain networks using graph metrics and has achieved great success in explaining the functional abnormality from the network mechanism[18]. However, this family of methods lack accuracy in the prediction task due to the model-driven methodology. Graph attention networks (GAT) [21], are novel neural network architectures that have been successfully applied to tackle problems such as graph embedding and classification. Different from CNN-based neurodisorders interpretation [11], one of the benefits of attention mechanisms is that they allow for dealing with variable-sized inputs, focusing on the most relevant parts of the input to make decisions, which can then be used for interpreting the salient input features. Motivated by this, we propose an innovative Edge-weighted Graph Attention Network (EGAT) with Dense Hierarchical Pooling (DHP), where the underling graphs are constructed from the functional connectivity matrices and the node features consist of both the anatomical features and the statistics of the nodal connectivity. Our contribution is summarized as follows:

- We propose a novel multi-modality analysis framework combining the sMRI and fMRI imaging in a graph classification task with workable settings.
- Our model outperforms the existing methods with an 10 ∼20% improvement, showing the necessity of multi-modality and attention infrastructures.
- We provide an interpretable visualization to understand the co-activation pattern of sMRI and fMRI from their activation maps.

## 2 Methodology

### 2.1 Construction of Graphs

On a labeled graph set *𝒞* = {(*G*_1_, *y*_1_), (*G*_2_, *y*_2_), …}, the general graph classification problem is to learn a classifier that maps *G*_*i*_ to its label *y*_*i*_. In practise, the *G*_*i*_ is usually given as a triple *G* = (*V, E, X*) where *V* = {*v*_1_, … *v*_*N*_} is the set of *N* nodes, *E* = {*e*_*ij*_}_*N*×*N*_ is the set of edges with *e*_*ij*_ denoting the edge weight, and *X* ∈ ℝ^*N*×*F*^ is the set of node features.

In our BP vs. HC binary classification setting, the nodes are defined by the region of interest (ROI) from some given atlas. For the edges, we utilize the densely connected graph rather than setting a threshold that dismisses the weak connectivity. The edge weight is then defined as the correlation-induced similarity given by 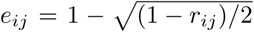, where *r*_*ij*_ is the Pearson’s correlation between the region-averaged BOLD time-series for region *i* and *j*. For each node, we construct a dim-11 feature vector combining the structural and functional MRI. The seven anatomical features are *Number of Vertices (NumVert), Surface Area (SurfArea), Gray Matter Volume (GrayVol), Average Thickness (ThickAvg), Thickness Standard Deviation (ThickStd), Integrated Rectified Mean Curvature (MeanCurv)* and *Integrated Rectified Gaussian Curvature (GausCurv)* [7], which provide the geometric information of brain surface. The four functional features are from connectivity statistics: mean, standard deviation, kurtosis and skewness of the node’s connectivity vector to all the other nodes, which summarize the moments of the regional time-series.

### 2.2 Graph Neural Network (GNN) Classifier

The architecture of our proposed GNN network is shown in Figure 1. Each graph *G* is first fed to a 5-heads EGAT layer, followed by two pooling layers that coarsens 129 nodes to 32/16 then to 4 for graph feature embedding. The extracted features are then fed to 2 fully-connected layers for classification.

**Fig. 1:**
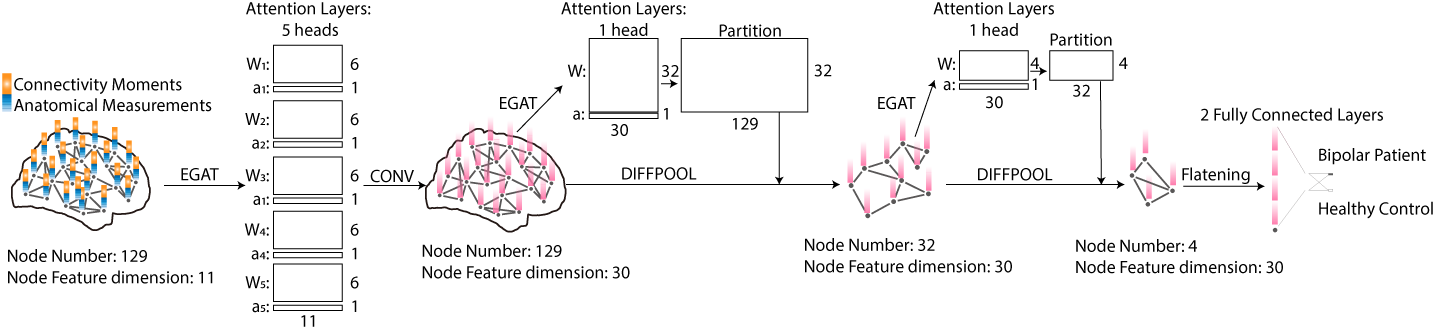
Schemata of EGAT-DHP classification network.

#### Edge-weighted Graph Attention Layer (EGAT)

The Graph Attention Layer takes a set of node features *X* = {***x***_1_, ***x***_2_…***x***_*N*_}, ***x***_***i***_ ∈ ℝ^*F*^ as input, and maps them to ***Z*** = {***z***_1_, ***z***_2_…***z***_*N*_}, ***z***_***i***_ ∈ ℝ^*F*^′. The idea is to compute an embedded representation of each node *v* ∈*V*, by aggregating its 1-hop neighborhood nodes {***x***_*j*_, ∀*j* ∈*N*(***x***_*i*_)} following a self-attention mechanism `Att`: ℝ^*F*^′ ×ℝ^*F*^′ → ℝ [21]. Different from the original [21], we leverage the edge weights of the underlying graph. The modified **attention map** *α* ∈ ℝ^*N*×*N*×*P*^ can be expressed as a single feed-forward layer of ***x***_*i*_ and ***x***_*j*_ with *edge weight e*_*ij*_:

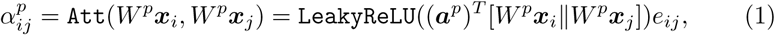

where *α*^*p*^ is the attention weight for the *p*-th attention head and 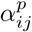 indicates the importance of node *j*’s features to node *i* in head *p*. It allows every node to attend all the other nodes on the graph based on their node features, weighted by the underlying connectivity. The *W*^*p*^ ∈ ℝ^*F*^′^×*F*^ is a learnable linear transformation that maps each node’s feature vector from dimension *F* to the embedded dimension *F*′. With *P* attention heads, attention mechanism `Att` is implemented by a **nodal attributes learning vector *a***^*p*^ ∈ ℝ^2*F*^′and `LeakyRelu` wit nput slope = 0.2. Then, the aggregation operation is defined as 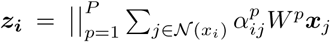 symbol ∥ represents the concatenation operation.

#### Dense Hierarchical Pooling (DHP)

To aggregate the information across nodes for graph level classification, we incorporate Dense hierarchical Pooling (DHP [23]) to reduce the number of nodes passing to the next layer. At the last level, the graph nodes are reduced to a few and features are flatten to a single vector, which is then passed to MLPs to generate graph label. The pooling procedure is performed by an assignment matrix ***S*** ∈ℝ^*N*^×^*N*^′ that coarsens both the node and edge information: ***z***_*out*_ = ***S***^*T*^ ***z***_*in*_, ***E***_*out*_ = ***S***^*T*^ ***E***_*in*_***S*** to a graph of *N*′ nodes. The assignment ***S*** is learned through another EGAT layer with the regularization loss *L*_*reg*_ = ∥***E, SS***^*T*^ ∥_*F*_, where ∥ · ∥_*F*_ denotes the Frobenius norm.

#### Neurological Motivation of Network Designing

Compared to the GCNs [10] with spectral convolution, our proposed GNN architecture allows for better description of local integration of node features, which is more biologically consistent with the findings of community structure of brain networks [14]. Secondly, the efficiency of hierarchical pooling lays on the implicit assumption that the underlined graph possesses the inferred structure. Thus, considering the typical numbers of communities discovered in previous literature [16] and the fact that the brain consists of four lobes, we add two pooling layers in our network where the first one pools the node set into 16/32 clusters and the second one pools the node set into 4 clusters. In addition, considering the heterogeneity of the brain networks in local signal processing, multiple heads are employed in the first layer of EGAT convolution.

### 2.3 Interpretation From Attention Map

Characterizing BP from anatomical MRI and task-fMRI and interpreting the brain features captured by the proposed model can help neuroscientists better understand BP. The **attention map** *α* in the EGAT layer learns salient cerebral cortex functional connectivity to identify BP by stacking layers in which nodes are able to attend over their neighborhoods’ features. By exploring the **gradient sensitivity** 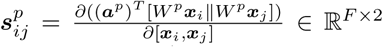, we can disentangle the relationship among node features (from different modalities) in identifying BP by examining the co-activation.

## 3 Experiment And Results

### 3.1 Image acquisition and processing

Data for this study consisted of 106 subjects (59 patients, 47 health controls) with 2 paired scans over 6 months. Both structural T1 MR (sMRI, dimension 192 × 256 × 256, voxel size 1 × 1 × 1*mm*^3^, fov= 192mm) and the functional MR (BOLD, dimension 64 × 64 × 30 × 244, voxel size 4 × 4 × 5*mm*^3^, fov 256, TR=3 s) scan were acquired on a GE 3-T scanner. During the fMRI scans, subjects performed “N-back” task in a block design manner (30 s/block, 11 blocks in total). We ended in 150 sMRI and fMRI pairs (75 patients, 75 health controls) after removing high-motion data (≥ 0.2 relative mean). Data was split into 5 folds based on subjects for cross-validation.

FreeSurfer [7] version 6.0 was employed in the image preprocessing for sMRI and the extraction of anatomical statistics. Image preprocessing for fMRI was done using FEAT pipeline of FSL [9] version 6.0, including steps of motion correction, spatial smoothing (FWHM 5), and registration to standard NMI space. A 0.01Hz high-pass filter was applied. We extracted regional mean BOLD time series with the *N* = 129 region in Lausanne atlas [5] and calculated the edge weights, connectivity matrices and functional features as described in Section 2.1. The functional connectivity matrices was then used as the underlined graph for EGAT. We also normalized each node feature separately by z-scores manner considering the heterogeneity for different measurements.

### 3.2 BP v.s. Healthy Control Classification

We investigated the best EGAT architecture by tuning the number of kernels. The experiment was run on 8 GTX Titan Xp (batch size=8) with Adam optimizer(learning rate=0.0001, betas=(0.9, 0.999), eps=1e-8). The optimal solution was achieved when the first pooling layer output 32 communities and the fully connected layer consisted of 32 nodes (see Table 1). The accuracy varied yet not too much when we changed the community size to 16 and the number of nodes in the FC-layer.

**Table 1:** Classification performance of different models (mean(std)%)

| model | accuracy | f1-score | precision | recall |
| --- | --- | --- | --- | --- |
| EGAT DiffPool(32-4) 32FC (*) | **82.00(3.80)** | **82.76(3.80)** | 79.46(4.45) | **86.67(6.67)** |
| EGAT DiffPool(16-4) 32FC | 81.33(5.05) | 80.84(5.95) | **82.38(4.26)** | 80.00(10.54) |
| EGAT DiffPool(32-4) 64FC | 80.67(2.79) | 81.39(2.67) | 80.27(11.61) | 85.33(12.82) |
| EGAT DiffPool(32-4) 16FC | 80.00(4.71) | 79.67(5.79) | 80.31(4.22) | 80.00(11.15) |
| GraphSAGE DiffPool(32-4) 32FC | 69.33(7.96) | 66.43(14.46) | 70.74(4.56) | 66.67(25.39) |
| MLP 32FC | 72.66(1.49) | 73.11(3.50) | 73.19(7.68) | 76.00(3.13) |
| Random Forest | 62.00(5.81) | 61.62(5.43) | 63.19(7.26) | 61.33(8.84) |
| Linear SVM | 58.67(8.84) | 59.84(7.55) | 60.20(10.13) | 62.67(16.11) |
| Our Model * (only fMRI) | 70.67(2.79) | 71.05(5.16) | 70.08(3.58) | 73.73(13.03) |
| MLP 32FC (only sMRI) | 68.04(5.66) | 67.71(12.24) | 68.35(7.74) | 73.21(23.42) |

To illustrate the importance of integrating multi-modality data, we compared the performance of using single modality. The results were shown in Table 1. First, to show the necessity of including anatomical features, we replaced the anatomical features as dummy variable ones (namely fMRI only) and performed the task with the same infrastructure as EGAT. The accuracy and F1-score decreased to 70.67±2.29% and 71.05±5.16% correspondingly. The results suggested that the anatomical features provided additional information. For the necessity of functional connectivity, we adopted a 2-layer MLP to classify the two groups based on the vectorized anatomical features of all regions (namely sMRI only). The accuracy and F1-score decreased to 68.04 ± 5.66% and 67.71 ± 12.24% correspondingly. The decreased performance showed the advantage of combining functional data in our proposed model.

To prove that our model better embedded both structural and functional features, we compared the accuracy and F1-score of our model versus Random Forest, SVM with Linear kernel and GraphSAGE (best parameters chosen by grid search). See Table 1, our model improved 10 ∼ 20% in the accuracy and F1-score, comparing with the three alternative models. The improvement may come from two causes. First, due to intrinsic complexity of sMRI and fMRI, complex models with more parameters is desired, which also explained why the MLP performed better than the other two. Second, as explained above, our model utilized the specific topology of the community structure in the brain network thus potentially modeled the local integration more effectively. We reached the conclusion that, not only the integration of multi-modality was necessary for imaging-based classification, but the proper modeling of the cross-modality also made a difference in disentangling the underlined complexity.

### 3.3 Biomarkers Discovery from Structural and Functional Features

One obstacle of applying complex models in diagnosis is the lack of interpretation. Here we utilize activation map and gradient sensitivity to show that our method can provide interpretable visualization of effective features on both group and individual levels in addition to the better prediction accuracy shown above. First of all, in panel a) of Figure 2, we showed the reordered attention map of each head averaged on all subjects. The chord diagram displayed the location and weight of edge-attention. We assigned colors to different brain regions and labeled their name at the bottom of panel a) of Figure 2. Second, in panel b) of Figure 2, we presented the gradient sensitivity of different node features. The gradient sensitivity on the node feature displayed two modes, one having weights on the source and target nodes with opposite signs and the other with same signs. We can see that the activation patterns are spatially selective, suggesting that the abnormality of biomarkers happened in a heterogeneous way on the brain network, except for Attention 4 that gave a quantification of the overall effect.

**Fig. 2:**
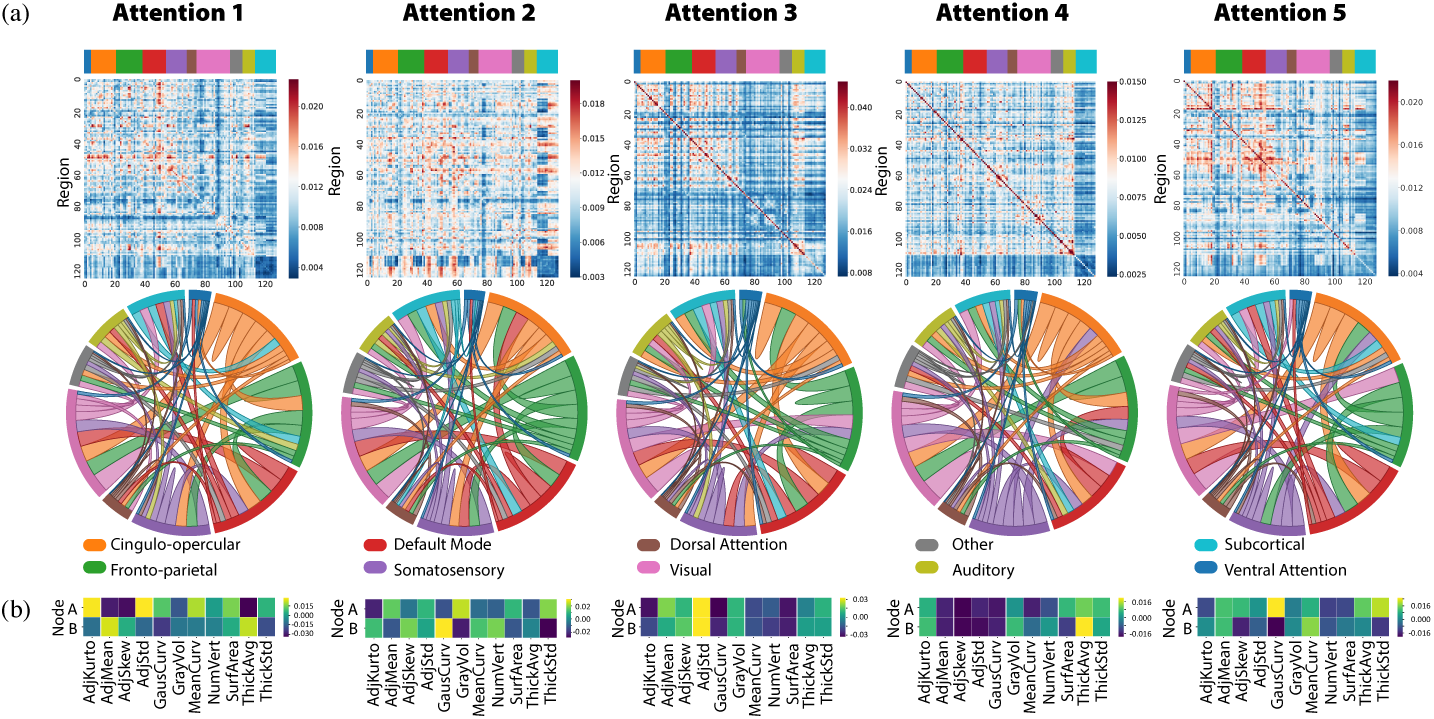
a) Activation maps and b) node feature gradient sensitivity of the five attention heads.

Attention 1 and 3 placed strong weight on the connectivity statistics in the node features with opposite modes. This indicated that these two attentions emphasized the heterogeneity of functional connectivity in two modes, mean of variance for Attention 3 and variance of variance for Attention 1. Combined with the spatial preference on the default mode network (DMN), fronto-parietal (FP) and cingulo-opercular (CO) networks, this supports the previous finding on the increase of regional homogeneity in the BD patients [13] and suggests potential sub-types in this deficit. While the focus on DMN in Attention 1 suggested that the integration and segregation of DMN could play a central role in psychiatry [15], the strong co-activation of connectivity and anatomical measurements suggested that the abnormality for DMN, FP and CO in functional networks could be associated with the deficit of anatomical properties [12,20].

For Attention 2 and Attention 5, the highest node weight was on the Gaussian curvature and complemented each other on the sign. Gray matter volume and thickness were also emphasized in these two attentions. While previous literature found widespread of gray matter deficit [12,20] but not atrophy in the white matter, our results here suggest that the white matter abnormality might be better represented by the curvature information [6]. Also, the spatial highlight on the cingulo-opercular (CO) besides the DMN supports the hypothesis that the deficit of CO integrity could be a reason of the deficit of cognition [17].

## 4 Conclusion

In this work, we proposed a novel graph-attention based method for cerebral cortex analysis that integrates sMRI and fMRI using GNN to classify BP v.s. HC. It helps to identify the unique and shared variance associated with each imaging modality that underlies cognitive functioning in healthy controls and impairment in BP. Thus, our model shows an superiority over alternative graph learning and machine learning classification models by 10 ∼ 20% in the accuracy and F1-score. In addition, by investigating the attention mechanism, we show that the proposed method not only provides spatial information supporting previous findings in the network-based analyses but also suggested a potential association of anatomical deficit and the abnormality of the functional network. This method can be generalized on multi-modality learning on neuroimaging.

